# Developing selective FPR2 agonists can be a potential approach to treat moderate to severe asthma

**DOI:** 10.1101/2021.09.16.460577

**Authors:** Senthil A. Visaga, Harikesh Kalonia, Vinay Verma, Sandeep Sinha, Shashi Kant Singh, Swati Upadhyay, Sudhir Sahdev, Amita Pansari, Rajesh Kumar, Mahadev Bandgar, Narayan Karanjule, Raj Kumar Shirumalla, Kaoru Morishita, Ruchi Tandon

## Abstract

Formyl peptide receptor (FPR) family members have been reported to play important role in the resolution of inflammation. A few FPR2/FPR1 dual agonists are reported in the public domain for their anti-inflammatory properties. None of these molecules, however, have been successful as a therapy yet. Recent reports bring forward the ambiguous role of FPR1 in inflammation. These include both positive and negative outcomes. We, therefore, aimed to develop selective FPR2 agonists and evaluated their potential in mitigating the non-resolving inflammation in mouse models of moderate to severe asthma. Extensive structure-activity-relationship (SAR) studies were conducted on the imidazole and benzimidazole chemotype series to identify potent and selective FPR2 agonists. A few molecules were shortlisted based on their *in vitro* profile and absorption, distribution, metabolism and excretion (ADME) properties and were further evaluated in mouse models of asthma. We report herewith identification of 3 RCI compounds with low nanomolar potency for FPR2 agonism and >10,000 fold selectivity over FPR1 in Ca^2+^ release assay. These molecules also showed potency in other *in vitro* assays and potent efficacy in three distinct animal models of asthma. Our data suggest that FPR2 agonism can be a potential therapeutic approach to treat asthma. Our findings also propose that FPR1 can be spared to achieve the desired pharmacological activity.

## 1. INTRODUCTION

Resolution of inflammation is an emerging concept which leads to the termination of inflammatory reaction by specific pro-resolving mediators (Perreti et al., 2015; Gómez et al., 2013). Emerging data suggests that by enabling healing and repair, the concept of resolution can be applied in treating various inflammatory diseases such as asthma, rheumatoid arthritis, Alzheimer and cardiovascular diseases (Prevete et al., 2015; Bozinovski et al., 2013; Cash et al., 2014;Kao et al., 2014; Le et al., 2002; Romano et al., 2015). Activation of Formyl peptide receptors seems to be an important facilitator of this interesting phenomenon. Formyl peptide receptors (FPRs) are G-protein-coupled receptors with three different isoforms; FPR1, FPR2, and FPR3 expressed in humans (Lu et al., 1992; Ye et al., 2009). FPR1 and 2 although have been studied in detail, information about the role of FPR3 is not very clear.

An increasing number of both peptide and non-peptide FPR1/2 dual agonists are being reported in the literature with demonstrated anti-inflammatory and immunological properties (Kao et al., 2014; Boulay et al., 1990, Bürli et al., 2006; Vacchelli et al., 2020; Qin et al., 2017; Cattaneo et al., 2019; Cilibrizzi et al., 2013; Cilibrizzi et al., 2009; Cilibrizzi et al., 2012; Crocetti et al., 2013; Giovannoni et al., 2013; Vergelli et al., 2016; Vergelli et al., 2017; Migeotte et al., 2006; Odobasic et al., 2018), none of them however have reached clinics so far. We, therefore, first conducted a careful literature analysis to understand the biology of FPR family members and if there are any functional differences between them, followed by in-house efforts to identify molecules with a desirable biological profile.

Concerning inflammation, FPR1 and FPR2 have contrasting but distinct roles. In endotoxemia studies, when FPR1 and FPR2 knock out mice (FPR1^-/-^ and FPR2^-/-^ respectively) were exposed to aerosolized lipopolysaccharide, lack of FPR1 but not FPR2 significantly reduced neutrophil infiltration in bronchoalveolar lavage and the lung tissue both intravascular and interstitial. Lack of FPR1 but not FPR2 inhibited lung edema and histological lung tissue alterations (Gómez et al., 2013).

FPR2^-/-^ mice data showed a convincing role of FPR2 in inflammation in another set of studies also (Dufton et al., 2013). The role of FPR1, however, has been very ambiguous (Vacchelli et al., 2020). On one hand, FPR1^-/-^ mice are protected from cigarette smoke-induced lung emphysema with a reduced migration of neutrophils and macrophages following cigarette smoke exposure (Cardini et al., 2012) and they also do not develop pulmonary fibrosis in response to intratracheal bleomycin challenge (Leslie et al., 2020). In a study using FPR1^-/-^ mice also, it has been found that FPR1 mediates the neutrophil recruitment in the lung on endotoxin challenge leading to acute lung injury (Grommes et al., 2014) suggesting an approach to develop FPR1 antagonists. On the other hand, FPR1 has been found to have a contradicting role in host defense in lung tissue. In one set of studies, FPR1^-/-^ mice are protected against Yersinia pestis infection (Owusu et al., 2019), and in another study, these mice show reduced clearance of *Escherichia coli* (Zhang et al., 2020) and *Listeria monocytogenes* infections (Liu et al., 2012) leading to increased levels of mortality. This rules out a strategy to develop FPR1 antagonists for asthma/lung injury. Besides, high expression levels of FPR1 in various cancers (Snapkov et al., 2016), its immunogenic role in celiac disease by inducing neutrophil migration (Lammers et al., 2015), and protective roles in ischemia-reperfusion studies and ventricular remodeling using FPR1^-/-^ mice (Zhou et al., 2018) indicate the ambiguous role of FPR1 in various other disease conditions also.

Given the anti-inflammatory potential demonstrated by FPR 1/2 dual agonists, FPR2 ^-/-^ knock out mice data, and the ambiguous data from FPR1 specific studies, we decided to develop selective agonists of FPR2. Our specific objectives were: i) To identify potent and selective FPR2 agonists with acceptable ADME properties, ii) to understand their molecular mode of action in cell-based assays, and iii) to evaluate the potential shortlisted FPR2 agonists in moderate to severe mouse models of asthma.

We conducted an extensive SAR to identify potent agonists of FPR2 belonging to the imidazole and benzimidazole series and shortlisted three molecules; RCI-1701306005, RCI-1701309674, RCI-1701307669 (Fig.1), based on their detailed biological profiling and ADME properties. RCI molecules showed the potencies in the nM range for FPR2 agonism and >10,000 fold selectivity over FPR1 in Ca^2+^ release assay. Selective FPR2 agonists also showed a time-dependent increase in the phosphorylation of extracellular signal-regulated kinase-1/2(Erk1/2) in CHO-FPR2 cells, as reported in some other studies (Qin et al., 2017, Cattaneo et al., 2019; Smirnova et al., 2010). Previously, cyclopentenone 15-deoxy--Δ^12,14^ -prostaglandin J2 (15d-PGJ_2_) has been reported to be a mediator of resolution of inflammation by activating MEK/ERK pathway (Cash et al., 2014, Marcone et al., 2016; Oliva et al., 2003; Takeda et al., 2001). We, therefore, tested the involvement of 15d-PGJ2 in FPR2 agonists induced ERK1/2 phosphorylation by RCI compounds. We found that the selective FPR2 agonists at our end increased the levels of 15d-PGJ2 in CHO-FPR2 cells confirming the molecular mechanism of resolution by these molecules. Asthma being a state of severe inflammation, we tried to explore the FPR2 agonist-mediated resolution in mitigating the same using three distinct mouse models of asthma. Interestingly, selective RCI molecules showed potent efficacy in all three progressive moderate to severe models of asthma. We, therefore, recommend that by promoting the resolution of inflammation, FPR2 agonists can be a potential therapeutic approach for asthma. Our findings also propose that FPR1 can be spared to achieve the desired pharmacological activity.

**Figure-1.**
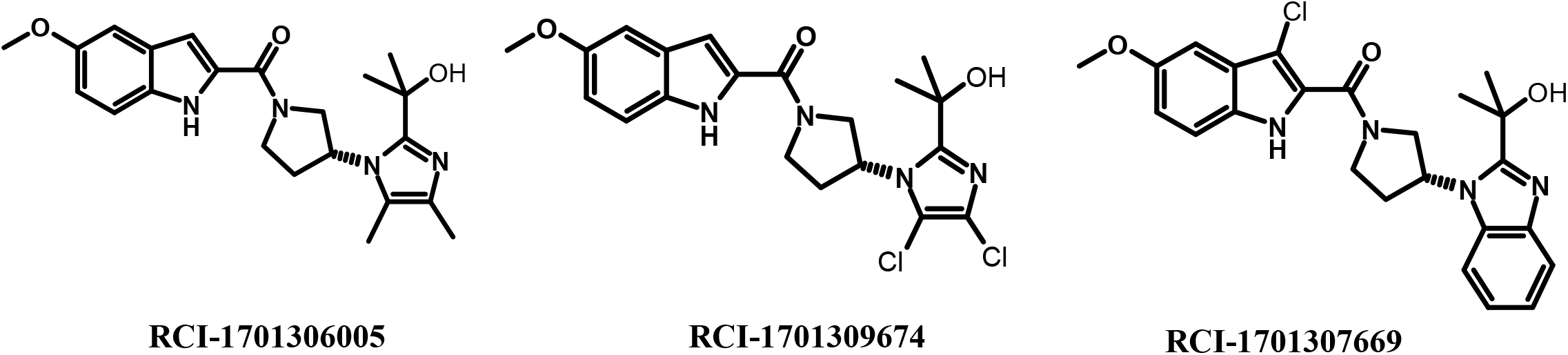
Chemical structure of selected compounds

## 2. MATERIALS AND METHODS

### 2.1 MATERIALS

#### 2.1.1 Cell culture and reagents

CHO cell lines overexpressing FPR2 and FPR1 receptors were generated in-house and propagated in Ham’s F12 medium (Gibco Biosciences) supplemented with 10% (v/v) heat-inactivated Foetal Bovine Serum (FBS) (Lonza Group Ltd.) containing 100 µg streptomycin ml^−1^ and 100 U penicillin ml^−1^ using neomycin (400 µg/ml) (Calbiochem, EMD Chemicals) as a selection marker. HL-60 Cells (ATCC CCL240) were procured from ATCC, Manassas, VA, USA. Cells were maintained in RPMI 1640 medium supplemented with 2 mM Glutamine and 10% (v/v) heat-inactivated FBS (Lonza Group Ltd.) containing 100µg streptomycin ml^−1^ and 100 U penicillin ml^−1^ and passaged as per the manufacturer’s instructions. Cells were maintained in a humidified atmosphere with 5% (v/v) CO_2_. 15d-PGJ 2 EIA kit was obtained from Enzo Life Sciences Inc., Farmingdale, NY, USA, and chemotaxis kit was obtained from Neuroprobe, Gaithersburg, MD, USA. All other chemicals used in the study were purchased from Sigma-Aldrich Corporation. Stock solutions of all the test compounds were prepared in dimethyl sulfoxide and further dilutions were made in KHB buffer (118 mM NaCl, 4.7 mM KCl, 1.2 mM MgSO_4_.7H_2_O, 1.2 mM KH_2_PO_4_, 4.2 mM NaHCO_3_, 11.7 mM D-Glucose, 1.3 mM CaCl_2_, and 10 mM HEPES) containing 30% DMSO. The final DMSO concentration in all the experiments was <0.5 %.

#### 2.1.2 Experimental Animals

All animal studies were conducted under the guidelines laid down by the Committee for Control and Supervision of Experiments on Animals (CPCSEA), Government of India, and were approved by the Institutional Animal Ethics Committee (IAEC) of Daiichi Sankyo India Pharma Pvt. Ltd. 7 weeks old female BALB/c mice were used for all the studies and were procured from Vivo Bio Tech Ltd., Telangana, India and were housed in individually ventilated cages in experimental rooms. Mice had access to irradiated standard pellet feed and filtered autoclaved water provided *ad libitum*. In each experimental group, 6-8 animals were taken.

### 2.2 METHODS

#### 2.2.1 Transfection of CHO-K1 cells

CHO-K1 stable cell lines over-expressing Homo sapiens FPR2 full length (NCBI accession number NM_001462.3), Homo sapiens FPR1 full length, Mus musculus (NCBI accession number NM_008039.2), FPR1 (NCBI accession number NM_002029.3) were generated using pcDNA3.1 vector flanked by CMV immediate early promoter and BGH poly-A tail and neomycin as the selection marker. CHO-K1 cells were grown in 6 well plates in Ham’s F12 medium (Gibco Biosciences) supplemented with 10% (v/v) heat-inactivated FBS (Lonza Group Ltd.) containing 100 µg streptomycin ml^−1^ and 100 U penicillin ml^−1^ and 5 mM L-glutamine. One day before transfection media was changed to OptiMEM (Gibco). Transfection of plasmid DNA was done using lipofectamine 2000 (Life Technology) following the manufacturer’s protocol. 48 hrs after transfection spent media was collected and fresh media supplemented with neomycin (400 µg/ml) was added. Two weeks after repeating the selection media colonies were picked up separately and analyzed for expression. The colony showing the best expression level was used for further assay.

#### 2.2 Ca^2+^ mobilization assay

The effect of test compounds on FPR2/FPR1 agonist-induced Ca^2+^ release was measured in this assay [38]. Briefly, the CHO cells over-expressing human FPR2 or FPR1 receptor were harvested and suspended in KHB buffer containing 0.25 mM sulfinpyrazone and 3 μM Fura-2AM dye and incubated at 37°C in a CO_2_ incubator for 30 min, followed by 10 min incubation at room temperature. Cells were then washed thrice with KHB buffer containing 0.5% BSA and plated at a density of 5×10^4^ cells/well in a 96-well black clear bottom plate.

To determine the agonistic activity, test compounds were added using the robotic arm of the fluorimetric imaging system; FlexStation 3 Multi-Mode Microplate Reader (Molecular devices LLC, CA). Fluorescence intensity in each well was measured at an excitation wavelength of 340 and 380 nm and an emission wavelength of 510 nm every 5 s for 240 s at room temperature after automated addition of compounds. The maximum change in fluorescence during the first 3 m, over baseline was used to determine a response. The agonistic response induced by 5 nM of W peptide in the case of FPR2 and 5 nM of fMLF in the case of FPR1 was considered as 100% to calculate the % agonistic activity of the test compounds. The final DMSO concentration in all the wells was 0.5%. The dose-response curve was generated using Prism 6 (GraphPad Software Inc., San Diego, CA, USA) and EC_50_ values were calculated using nonlinear regression analysis. Data was represented as an average of 3 independent experiments (n=3) with a % CV value was <10 % CV and Z’ value in the range of 0.9-1.

#### 2.2.3 Erk1/2 phosphorylation and FPR2 expression studies

For phosphorylation studies, CHO-FPR2 cells were incubated with FPR2 agonists for the desired time point. Cells were harvested using 1x Trypsin EDTA, washed with ice-cold PBS and treated with ice-cold lysis buffer containing 20mM Tris-HCl (pH 7.5), 150 mM NaCl, 1mM EDTA, 1 mM EGTA,1% Triton-X, 2.5 mM Sodium pyrophosphate,1mM β-glycerophosphate,1mM Na_2_VO_4_, 1 µg/ml Leupeptin and 1 mM PMSF. Lysed samples were centrifuge at 11,000 x g for 10 minutes at 4° C and supernatants were separated by SDS-PAGE, transferred to a nitrocellulose membrane, and immune-blotted with rabbit-anti-human P44/4 (Erk1/2) antibody. Expression of FPR2 was confirmed rinsed with ice-cold PBS and lysates were prepared as above. Human FPR2 antibodies (Novus Biologicals) for FPR2 expression analysis or and goat-anti-rabbit IgG HRP (Cell Signaling Technology) were used as primary and respective secondary antibodies. Proteins were visualized by the ECL system.

#### 2.2.4 Total RNA extraction and RT-PCR

Total RNA from CHO-FPR2 overexpressing cells or HL-60 cells was isolated using Trizol reagent (Invitrogen, Carlsbad, CA, USA) and purified using RNeasy Mini kits (Qiagen, Valencia, CA, USA). Total RNA (1–2 μg) was reverse-transcribed in a 25 µl reaction using Taqman reverse transcription reagents (Applied Biosystems, CA, USA). RT-PCR multiplex reaction was performed using gene-specific Taqman primers and probes (Applied Biosystems, CA, USA) and analyzed using QuantStudio 6 Flex (Applied Biosystems, CA, USA) Real-Time PCR System. 18s endogenous control mRNA. Relative quantitation (RQ) of mRNA expression was analyzed using QuantStudio™ Real-Time PCR Software v1.7.1 (Applied Biosystems, CA, USA).

#### 2.2.5 15d-PGJ2 release in CHO-FPR2 cells

CHO-FPR2 cells were plated in a 96 well plate at a density of 5 × 10^3^ cells per well. Cells were treated with desired concentrations of FPR2 agonists the next day for 16 h. 15d-PGJ2 levels in cell culture supernatants of CHO-FPR2 cells were analyzed using an enzyme immunoassay kit following the manufacturer’s instructions and (Enzo Life Sciences Inc., Farmingdale, NY, USA). The plate was read in a microplate reader at 405nm and 15d-PGJ2 levels were calculated compared to untreated control.

#### 2.2.6 Chemotaxis assay

Hl-60 cells were differentiated in the presence of 1% DMSO for 7 days. Differentiated HL-60 cells are expected to behave like neutrophils. Chemotaxis was analyzed in 96 well ChemoTx chemotaxis chambers (Neuroprobe, Gaithersburg, MD, USA). 2×10^6^ differentiated HL-60 cells were re-suspended in 30 µl HBSS containing 0.5 % (v/v) FBS in the upper chamber of the chemotaxis chambers and the test compounds were added to the lower wells of chemotaxis assembly. HL-60 from the upper well was allowed to migrate to the lower wells through the 5.0 μm polycarbonate filter for 60 min at 37° in a CO_2_ incubator. The number of migrated cells was determined by measuring ATP in the lysates of transmigrated cells using a luminescence-based assay (CellTiter-Glo; Promega, Madison, WI, USA) and readouts were taken in the FlexStation 3 Multi-Mode Microplate Reader (Molecular de- vices LLC, CA). Luminescence was converted to the % value after considering the luminescence obtained using the total number of HL-60 cells as 100 %.

#### 2.2.7 Ovalbumin-induced airway eosinophilia

Female BALB/c mice weighing 25-30 g were sensitized with 100 µl normal saline containing 2 mg of aluminum hydroxide and ovalbumin (OVA) at 100μg/mice, intra-peritoneal (i.p) from day 1 to day 14. Animals in the sham control group were administered 2 mg of aluminum hydroxide i.p.

Mice were challenged with OVA at (50 μg/mice) dissolved in 50 µl normal saline under anesthesia by intranasal route followed by administration of vehicle (0.5 % methylcellulose) or test compound by the oral route on days 21 to 23. The sham-control group was challenged intranasally with 50 µg of ovalbumin dissolved in 50 µl of normal saline by the intranasal route. Vehicle/test compounds/reference standards were administered twice a day.

Mice were euthanized on day 23, with an overdose of sodium thiopentone administered intraperitoneally. Subsequently, broncho-alveolar lavage fluid (BALF) was collected by tracheostomy using ice-cold Hank’s Balanced Salt Solution (HBSS). The collected fluid was centrifuged at 2,000 x g for 10 min at 4°C and the cell pellet was re-suspended with 1 mL HBSS. The total leukocyte count in the cell suspension was enumerated by a veterinary hematology analyzer (Sysmex XT-2000iV, Sysmex, Japan). For obtaining differential leukocyte count, smears were prepared using 50 µl of the cell suspension on a frosted glass slide, stained with Leishman’s stain, and cell counts were taken by scientists who were blinded to the treatment groups.

#### 2.2.8 House dust mite (HDM) induced chronic airway inflammation and airway hyperresponsiveness

Mice of all groups except the sham control group were instilled *Dermatophagoides pteronyssinus* HDM extract (Greer Labs) intranasally (25 µg of protein in 20 µl normal saline) five days per week for five weeks. Sham-control animals received 20 µl of normal saline by the intranasal route. Vehicle/test compounds/reference standard was administered from week four till the last HDM exposure and was administered twice daily. Animals were administered the vehicle/test compounds/reference standard by the oral route.

Twenty-four hours after the last HDM instillation, mice were transferred to the body box of a whole-body plethysmograph (Buxco Laboratories). Subsequently, mice were exposed to the aerosolized saline following which the Penh was recorded for 5 min. Thereafter, the animals were sequentially exposed to aerosolized serotonin (Sigma-Aldrich) at the doses of 3, 10, and 20 mg/ml, and an increase in Penh was recorded for 10 min following each serotonin exposure. Airway hyperresponsiveness (AHR) score was computed as the sum of the broncho-constrictor responses to aerosolized serotonin. Mice were euthanized 24 h after recording the AHR and bronchoalveolar lavage fluid was collected and processed for the enumeration of total and differential leukocyte counts as described in section 2.8.

#### 2.2.9 HDM and Complete Freund’s Adjuvant (CFA) induced airway inflammation

On day 0, animals of all groups except the sham-control group were immunized subcutaneously on their backs with 100 µl of an emulsion containing 50 µl of Complete Freund’s Adjuvant (Sigma-Aldrich) and 100 µg of *Dermatophagoides pteronyssinus* HDM extract (Greer Labs; based on protein content) dissolved in 50µl of normal saline. Sham-control animals were administered an emulsion prepared with incomplete Freund’s adjuvant and normal saline.

On day 14, animals of all groups except the sham control group were intranasally challenged with 100 µg of HDM extract dissolved in 20µl of normal saline. Sham-control animals were administered normal saline by the intranasal route. One hour before the HDM-challenge, the animals were administered the vehicle/test compounds/reference standard by the oral route. Vehicle/test compounds/reference standard were administered twice a day and the dosing schedule was followed on day 15 as well. The animals were euthanized 48 h after HDM-challenge, bronchoalveolar lavage fluid was collected and processed for the quantification of total and differential leukocyte counts as described in section 2.2.8.

#### 2.2.10 LPS induced Neutrophilia

Female BALB/c mice (Body weight: 25-30 g) were administered orally with the vehicle (0.5 % methylcellulose) or test compound. One hour later, mice of all groups except the normal control group were challenged with LPS (25 μg/mouse administered intranasally). The normal control group was challenged with saline. The animals were euthanized 4 h after LPS/saline challenge, bronchoalveolar lavage fluid was collected and processed for the quantification of total and differential leukocyte counts as described in section 2.2.8.

#### 2.2.11 Statistical analysis

At least n=3 or more experiments were conducted and all the experimental conditions were run in triplicate in cell-based assays. S.E.M. was calculated for each data point. All the pooled data, therefore, was an outcome of 9 independent observations. Nonlinear regression analysis was used to determine the EC_50_ of the agonists using GraphPad Prism software. All animal experiments contained a minimum of 6-8 animals in each group. The significance of the data generated was confirmed by conducting Dunnet and student’s T test and a p-value of <0.05 were considered statistically significant.

## 3. RESULTS

### 3.1 Compounds RCI-1701306005, RCI-1701309674, RCI-1701307669 are potent and selective FPR2 agonists

Several selected compounds were evaluated in Ca^2+^ release assay using CHO cells over-expressing human and mouse FPR2. As a result of extensive structure-activity relationship studies, we identified RCI-1701306005, RCI-1701309674, RCI-1701307669 (Fig.1) with nM potency in Ca^2+^ release assay with no or little species variation using human and mouse FPR2 over-expressing cells (Fig.2, Table-1). Also, these compounds were found to be ≥10000 fold selective over hFPR1. We evaluated RCI compounds reported in this manuscript for other GPCRS also such as muscarinic 2 (M2), muscarinic 3 (M3), Urotensin I, Urotensin II receptors, and a panel of 42 kinases from various tyrosine and serine-threonine kinase families and these compounds did not show any liability @ 10 μM concentration.

**Figure-2.**
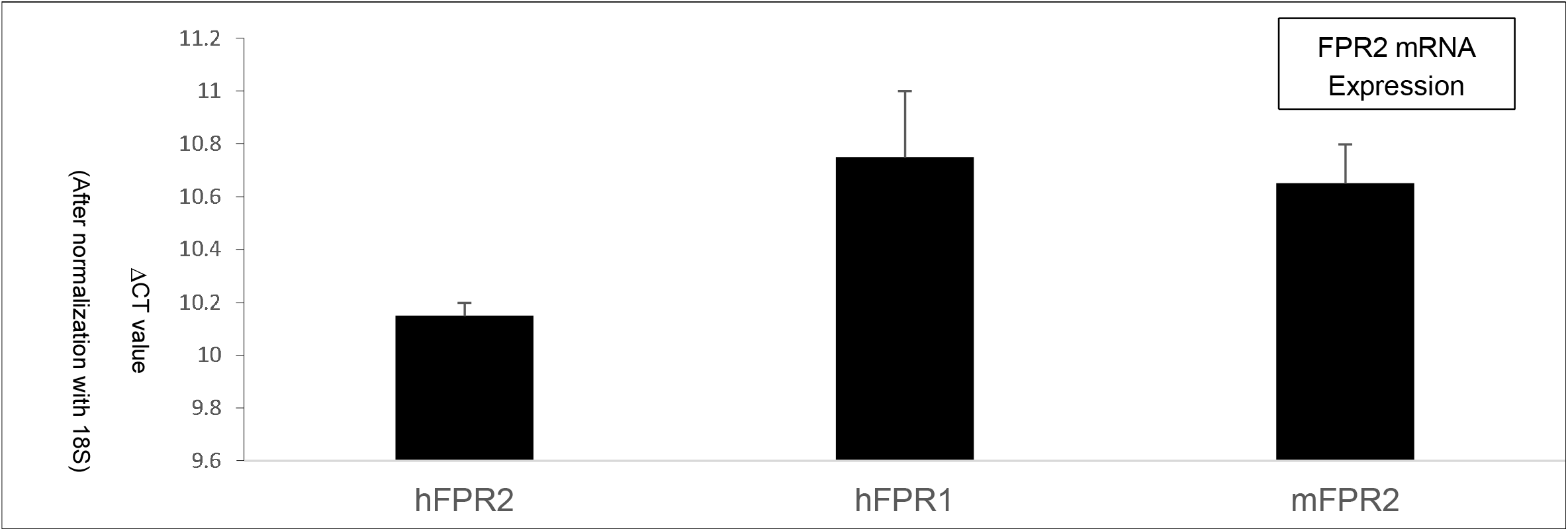
Expression level of FPR1 and FPR2 in CHO cells overexpressing FPR1 and FPR2 respectively by real time PCR

**Table-1.**
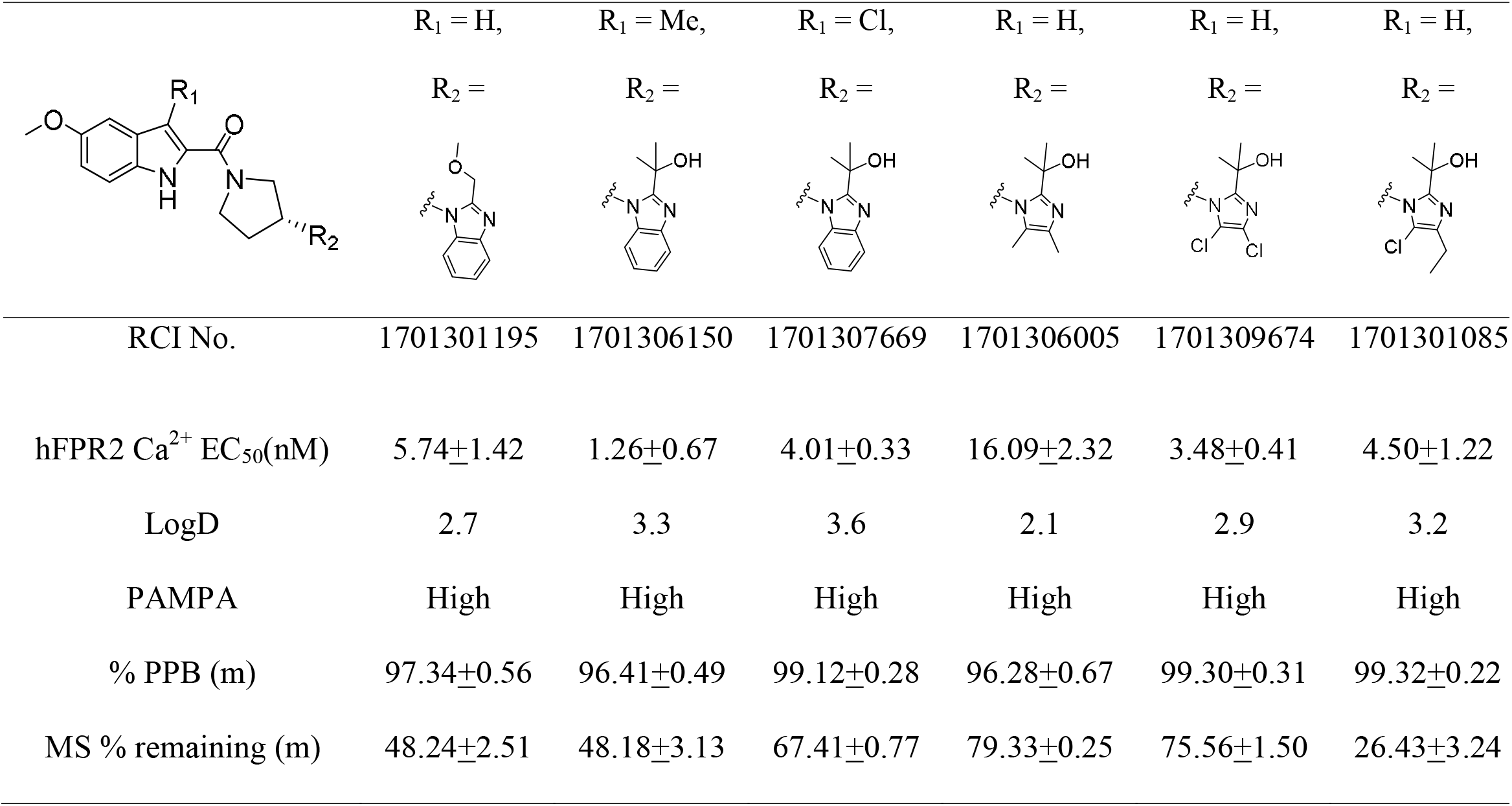
*In vitro* and ADME profile of selected NCEs (n=3)

|  | R<sub>1</sub> = H,<br>R<sub>2</sub> = | R<sub>1</sub> = Me,<br>R<sub>2</sub> = | R<sub>1</sub> = Cl,<br>R<sub>2</sub> = | R<sub>1</sub> = H,<br>R<sub>2</sub> = | R<sub>1</sub> = H,<br>R<sub>2</sub> = | R<sub>1</sub> = H,<br>R<sub>2</sub> = |
| --- | --- | --- | --- | --- | --- | --- |
| RCI No. | 1701301195 | 1701306150 | 1701307669 | 1701306005 | 1701309674 | 1701301085 |
| hFPR2 Ca<sup>2+</sup> EC<sub>50</sub>(nM) | 5.74±1.42 | 1.26±0.67 | 4.01±0.33 | 16.09±2.32 | 3.48±0.41 | 4.50±1.22 |
| LogD | 2.7 | 3.3 | 3.6 | 2.1 | 2.9 | 3.2 |
| PAMPA | High | High | High | High | High | High |
| % PPB (m) | 97.34±0.56 | 96.41±0.49 | 99.12±0.28 | 96.28±0.67 | 99.30±0.31 | 99.32±0.22 |
| MS % remaining (m) | 48.24±2.51 | 48.18±3.13 | 67.41±0.77 | 79.33±0.25 | 75.56±1.50 | 26.43±3.24 |

### 3.2 FPR2 agonists lead to the phosphorylation of Erk1/2 independent of PI3 kinase signaling cascade

Our results indicate that all the three shortlisted FPR2 agonists; RCI-1701306005, RCI-1701309674, RCI-1701307669 stimulated the phosphorylation of Erk1/2 in a time-dependent manner similar to W peptide, FPR1/2 dual agonist, using CHO cells over-expressing FPR2 receptor suggesting that FPR2 alone is sufficient to activate the pro-survival biomarkers. Since PI3 kinase has also been reported to regulate Erk1/2 signaling, we wanted to know whether the FPR2 agonist-mediated phosphorylation of Erk1/2 involves PI3kinase. We, therefore, evaluated the effect of PI3 Kinase inhibitor LY294002, on the phosphorylation of Erk1/2 by RCI compound as well as W peptide. Our data suggest that LY294002 did not affect the phosphorylation of Erk1/2 by these agonists. This suggests that FPR2 agonists activate Erk1/2 independent of PI3 kinase-mediated signaling cascade (Fig.3).

**Figure-3.**
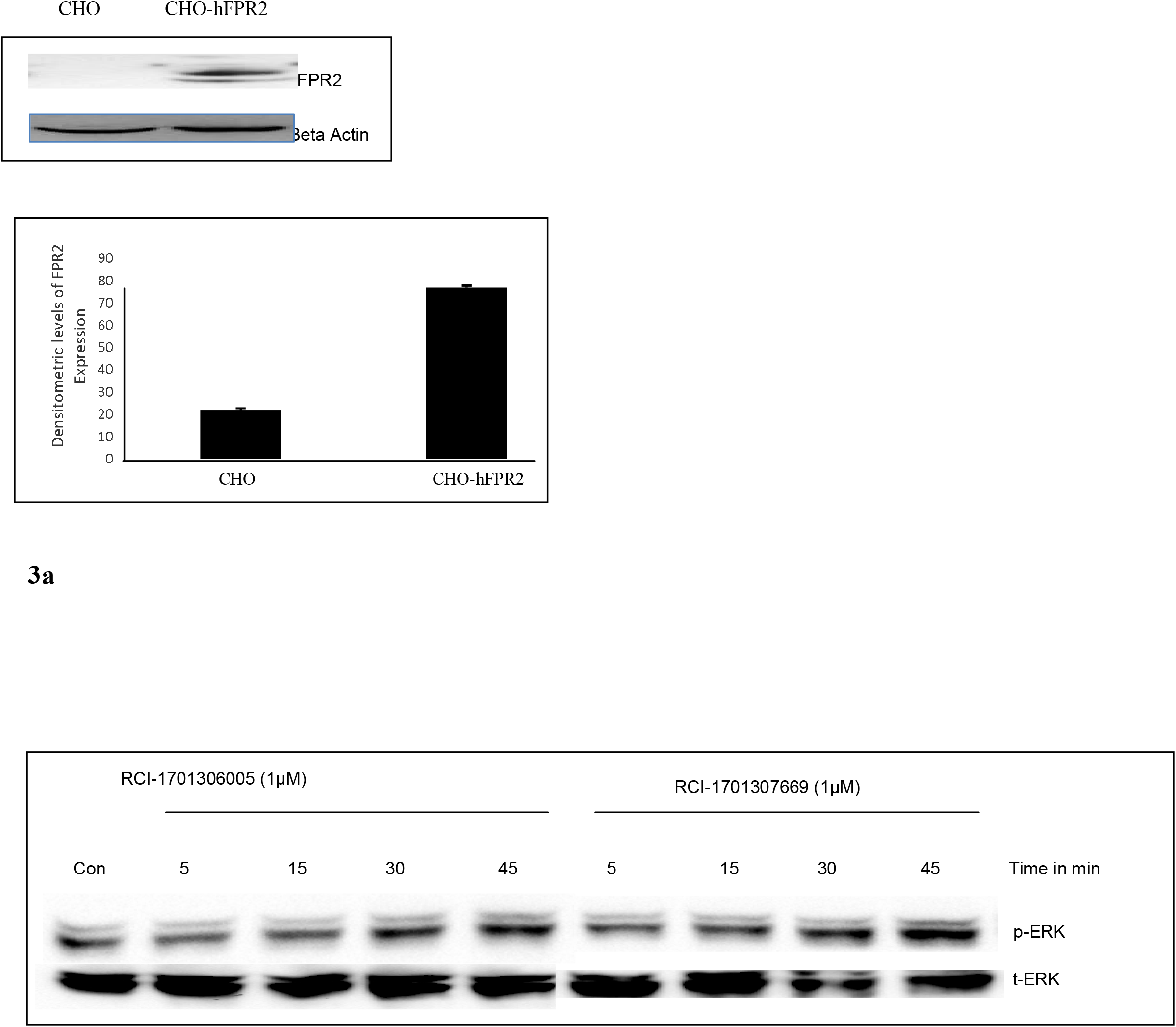
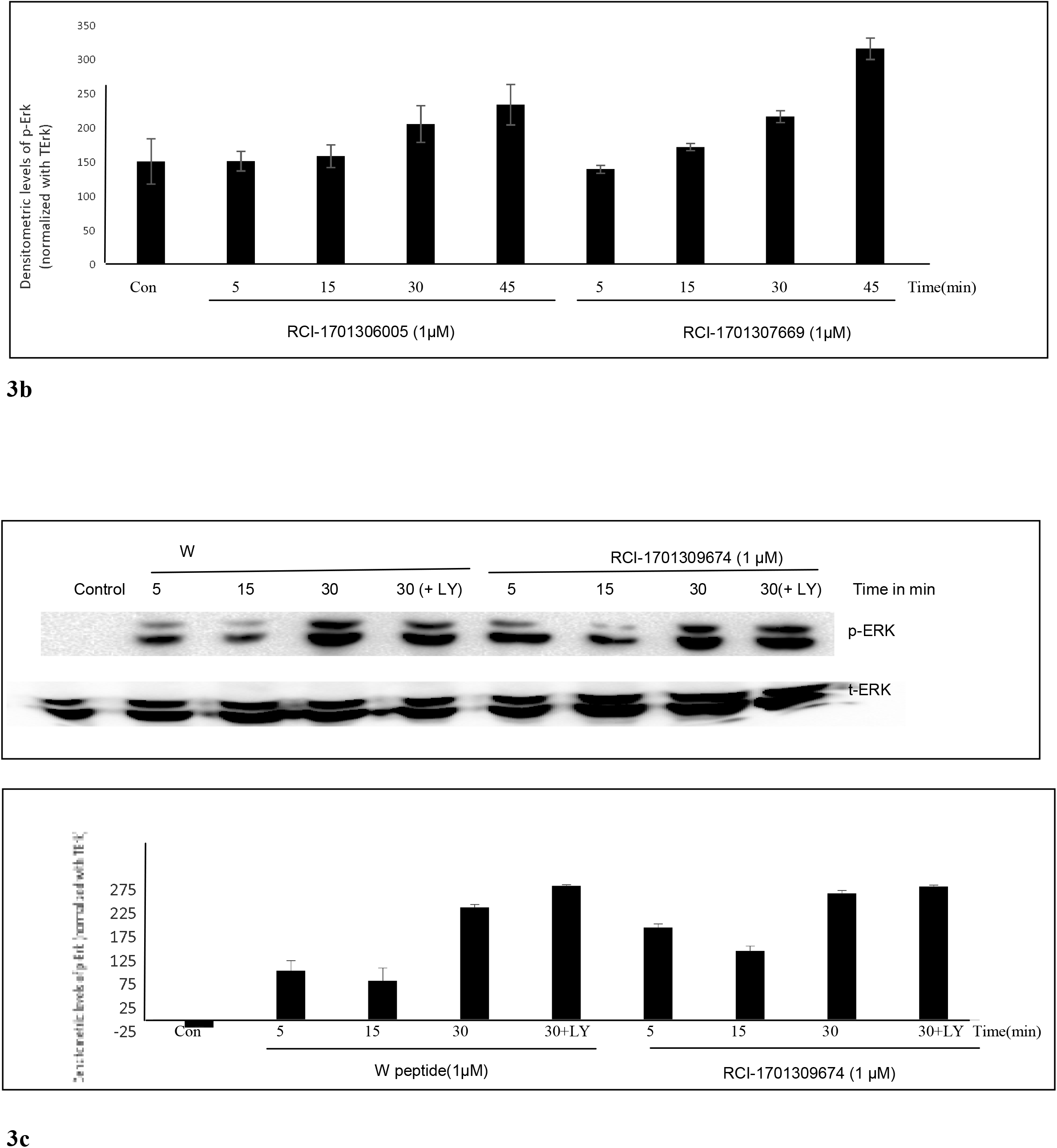
Erk1/2 phosphorylation by FPR_2_ agonists in CHO-FPR2 cells, as analyzed by western blot and their relative densitometric values after normalization with beta actin (for FPR2 expression analysis) or total Erk1/2 for phosphor Erk1/2 expression analyses): a) expression of FPR2 protein in CHO-hFPR2 cells, b) phosphorylation of Erk1/2 by RCI-1701306005 and RCI-170130669 and c) W peptide and RCI-1701309674 in a time dependent manner.

### 3.3 15d-PGJ2 release

Cyclopentenone 15-deoxy-**-**Δ^**12**,**14**^ -prostaglandin J2 (15d-PGJ2) has been previously reported to bind and activate H-Ras leading to phosphorylation and activation of Erk1/2 [5, 35-37]. We, therefore, tested the involvement of 15d-PGJ2 release by these compounds which may be linked to ERk1/2 phosphorylation. We found that all the three FPR2 agonists included in our study led to an increase in 15d-PGJ2 in CHO-FPR2 cells above basal levels in a concentration-dependent manner (Fig.4).

**Figure-4.**
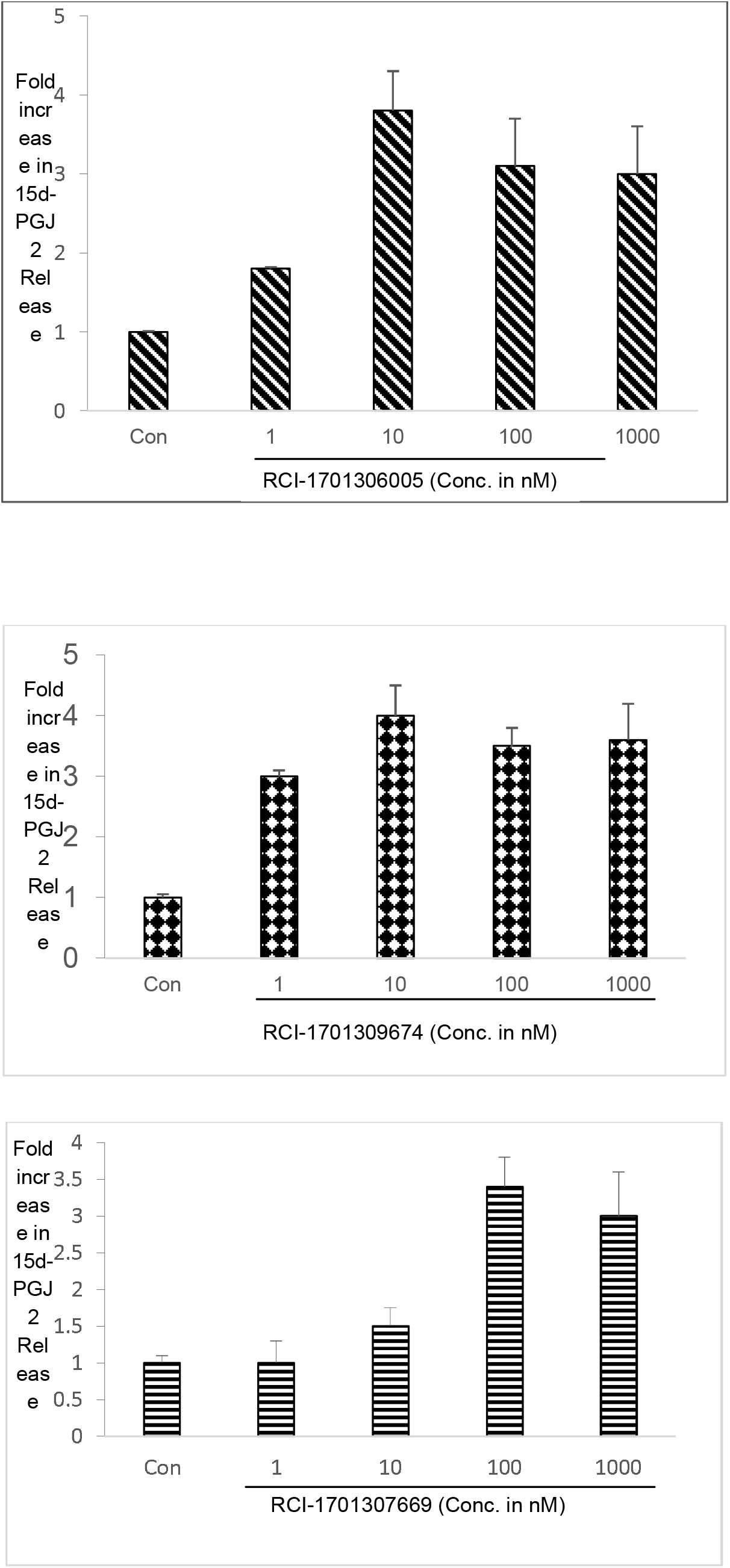
15dPGJ2 release in CHO-FPR2 cells to demonstrate Resolution mediated activity by RCI compounds

### 3.4 Chemotaxis

FPR2 is expressed by neutrophils and triggers chemotaxis. In absence of neutrophils from human blood, which requires approval from the human ethics protocol approval committee, we used differentiated HL-60 cells in our experiment as they have been previously reported to behave as neutrophil-like cells in response to dimethyl sulfoxide. Expression of FPR2 was confirmed in HL-60 cells, both pre and post differentiation by mRNA expression analysis using real-time PCR. Change in chemotaxis was ensured by FPR2 agonist using differentiated HL-60 cells compared to untreated controls (Fig.5).

**Figure-5.**
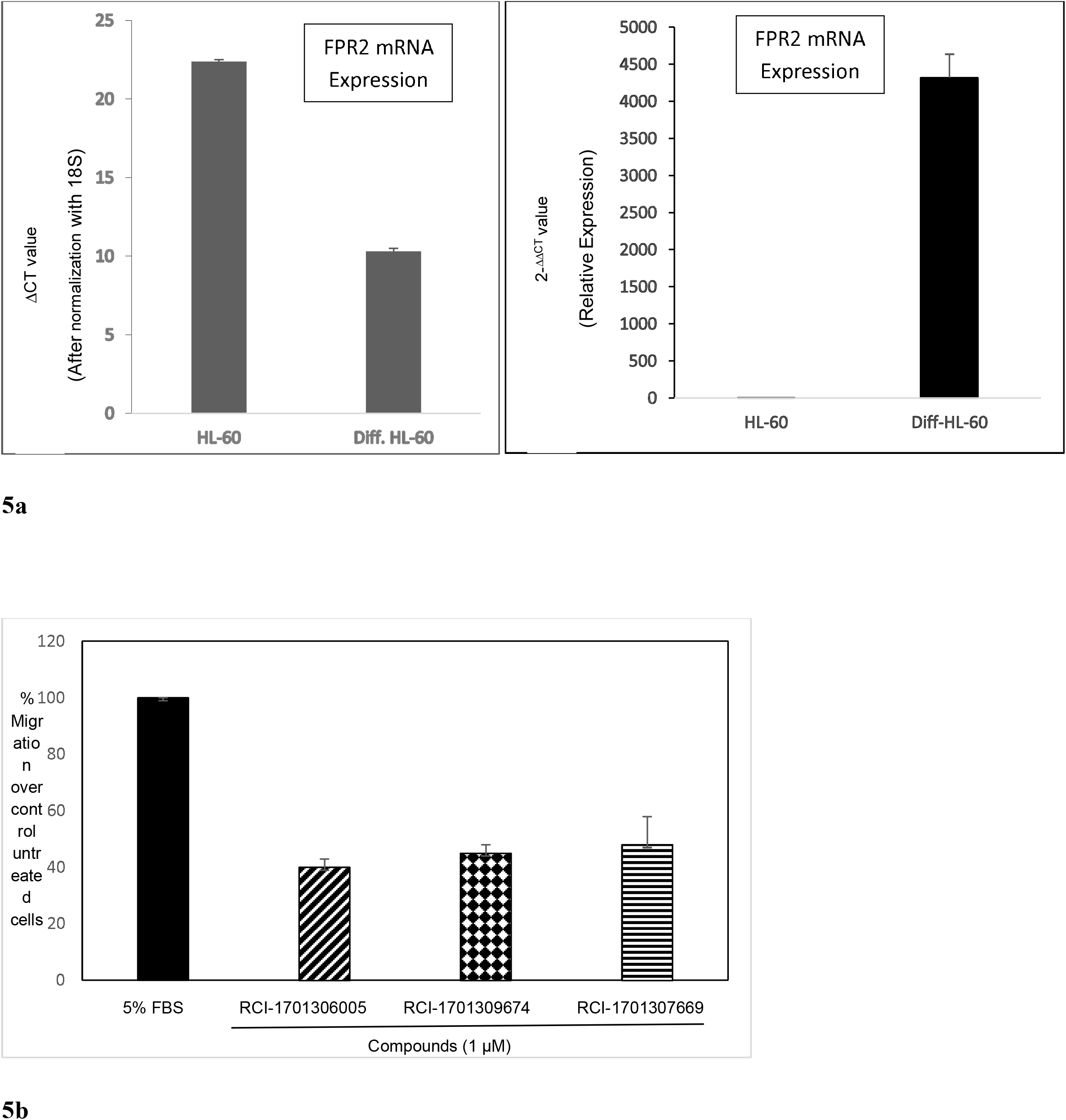
Chemotaxis Assay in HL-60 cell line a) real time PCR assay to suggest expression of FPR2 in differentiated and un-differentiated HL-60 cells b) % migration of differentiated HL-60 cells by RCI compounds

### 3.5 Pharmacokinetic analysis

Pharmacokinetic analysis was conducted to assess the exposure and bioavailability of these compounds at the tested doses. Oral PK was conducted at the dose of 10 and 100 mg/ in mice. PK study by intravenous route was conducted at 2mg/kg. Blood samples were collected after 1, 2, 4, 8, and 24 h of administration of test compound, and different pharmacokinetic (PK) features were analyzed (n =3), (Fig.6, Table-3). The dotted line suggests the EC_50_ values of these compounds in the cell-based assay and exposure of these compounds at the tested compounds was above the cell-based potency. Data are presented as mean ±SEM. The significance of the data generated was confirmed by conducting Dunnett and student’s T-test. A P-value of <0.05 was considered statistically significant.

**Figure-6.**
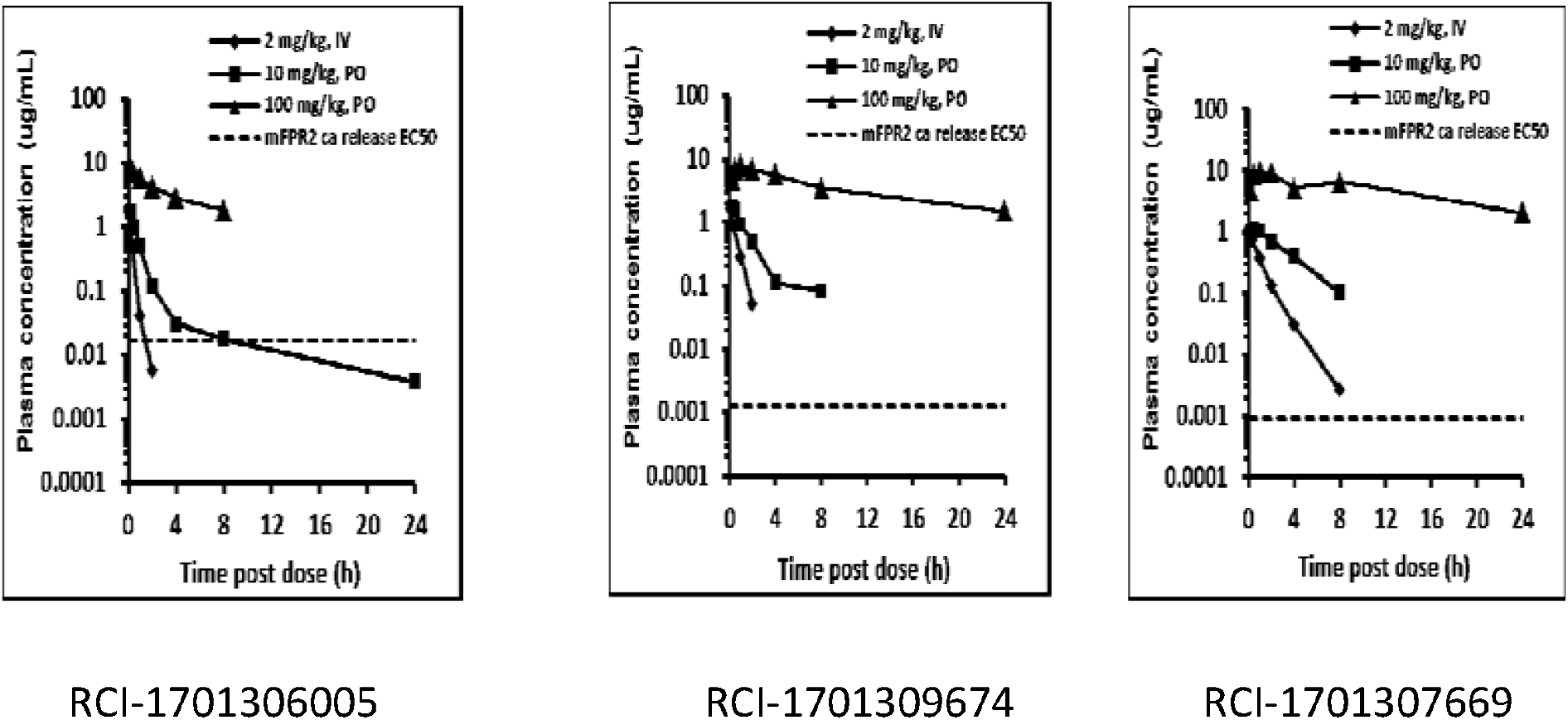
Pharmacokinetic profile of RCI compounds by i.v(2mg/kg) and p.o.(10 and 100 mg/kg) routes.

### 3.6 Efficacy of RCI compounds in mouse models of moderate to severe asthma

To assess the potential of FPR2 agonists for asthma, selected compounds were evaluated in the mouse OVA model. Ovalbumin challenge produced significant airway eosinophilia. Dexamethasone (1 mg/kg, p.o., q.d) which was used as a method of control, significantly inhibited airway eosinophilia and RCI compounds RCI-1701306005, RCI-1701309674, RCI-1701307669 also demonstrated dose-related inhibition of airway eosinophilia with ID_50_ values of 11.8, 10.7 and 14.7 mg/kg, b.i.d. p.o respectively(Fig.7). In an another severe model using house dust mice also, RCI compounds also showed significant inhibition of airway eosinophil influx in the HDM induced model of eosinophilia at 10 and 30 mg/kg, p.o., b.i.d. as well as inhibition of airway hyper-responsiveness at 30 mg/kg, p.o., b.i.d. (Fig.8) in Balb/c mice. In a progressively more severe model of HDM+CFA, these molecules inhibited the levels of eosinophils and neutrophil influx in the broncho-alveolar lavage (Fig.9). In LPS induced neutrophilia model also, FPR2 agonists showed inhibition of LPS induced neutrophilia in LPS model, as shown in Figure 10 (NEW).

**Figure-7.**
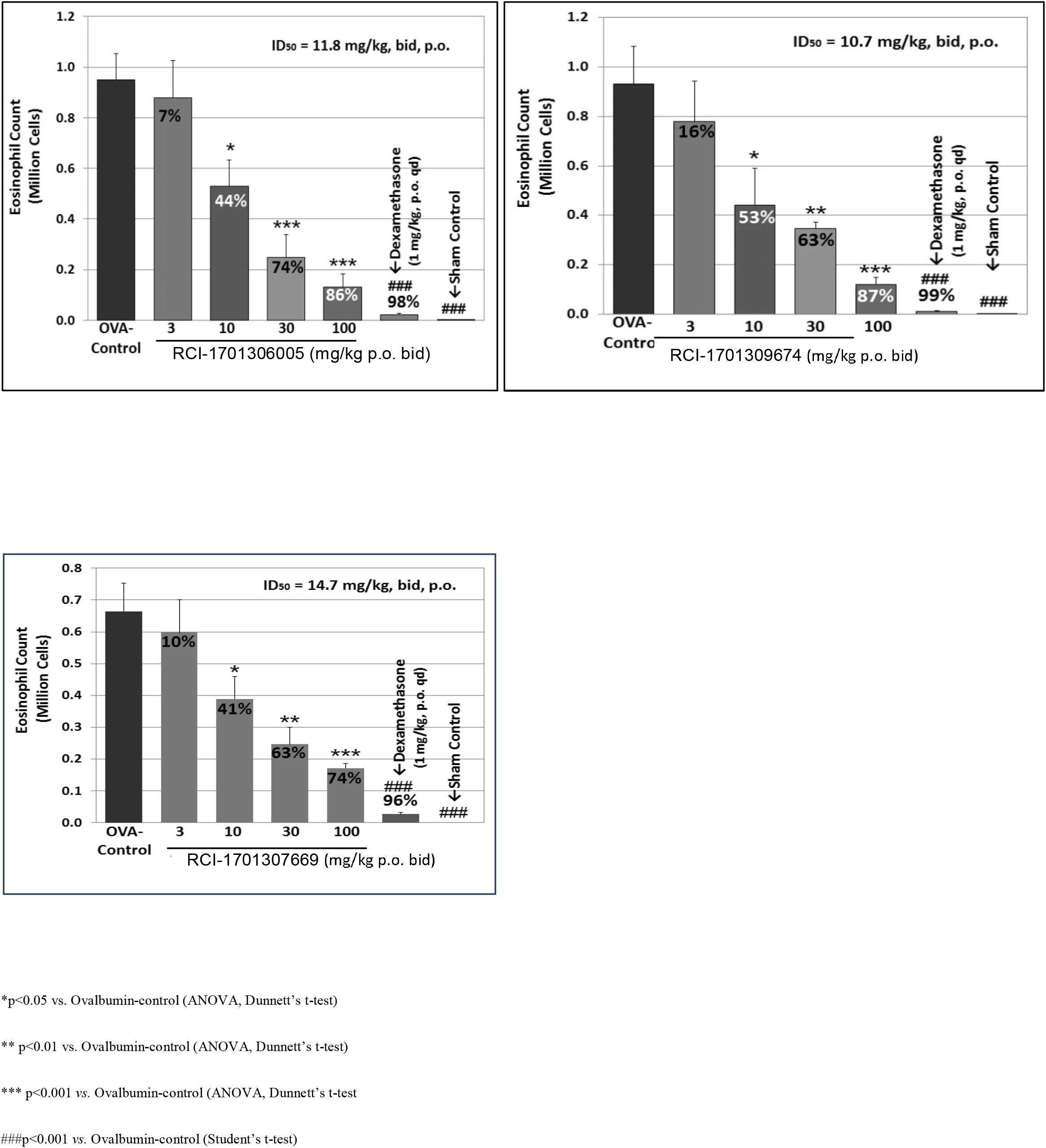
Dose response of RCI compounds in OVA induced airway eosinophilia mouse model in BALB/c mice tomonitor change in eosinophil counts

**Figure-8.**
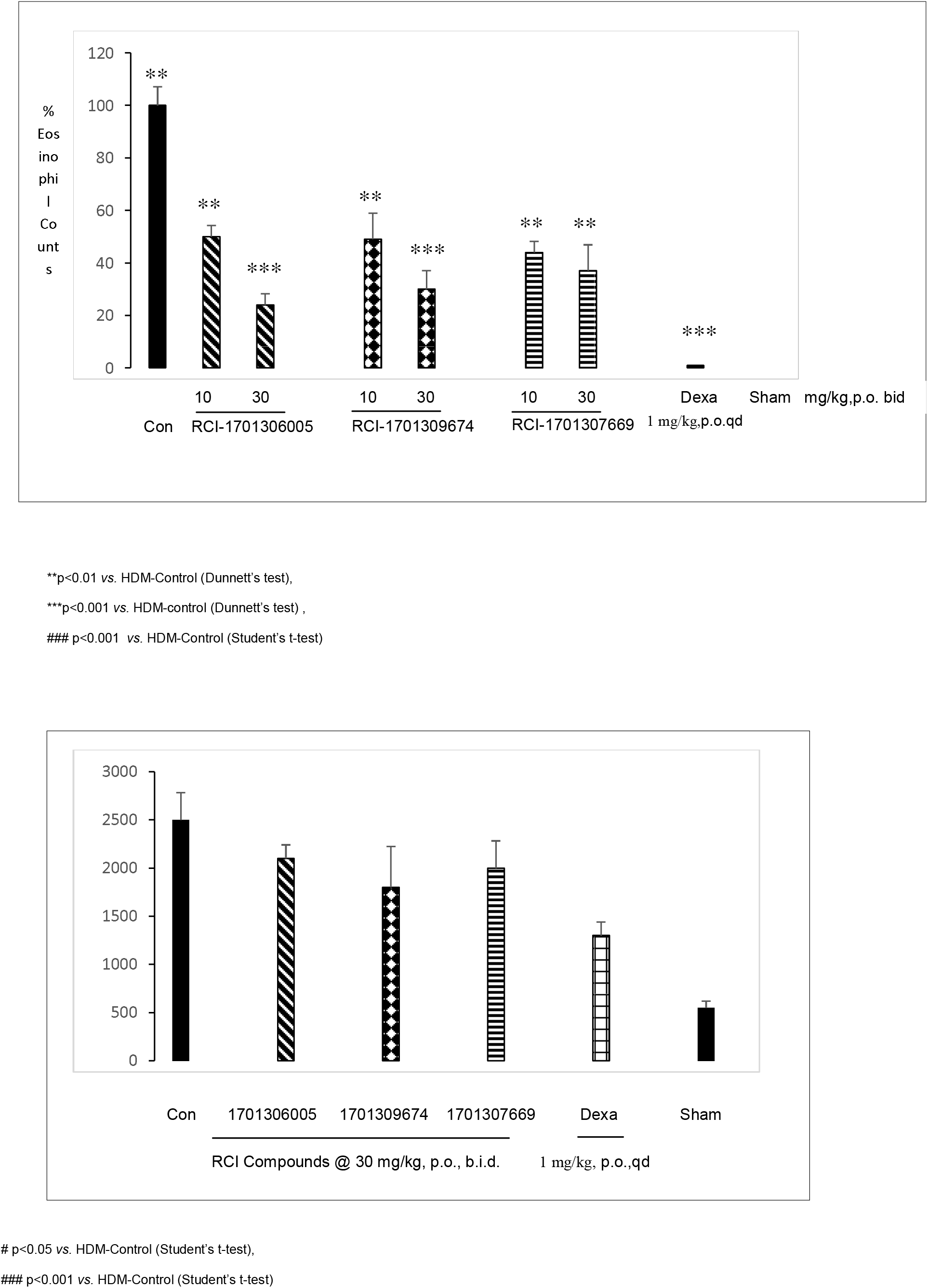
Evaluation of RCI compounds in HDM induced mouse model in BALB/c mice to monitor a)change in eosinophil counts by test compounds at 10 and 30 mg/kg p.o. dose to monitor change in eosinophil counts b)modulation of airway hyper responsiveness by RCI compounds at 30 mg/kg p.o.

**Figure-9.**
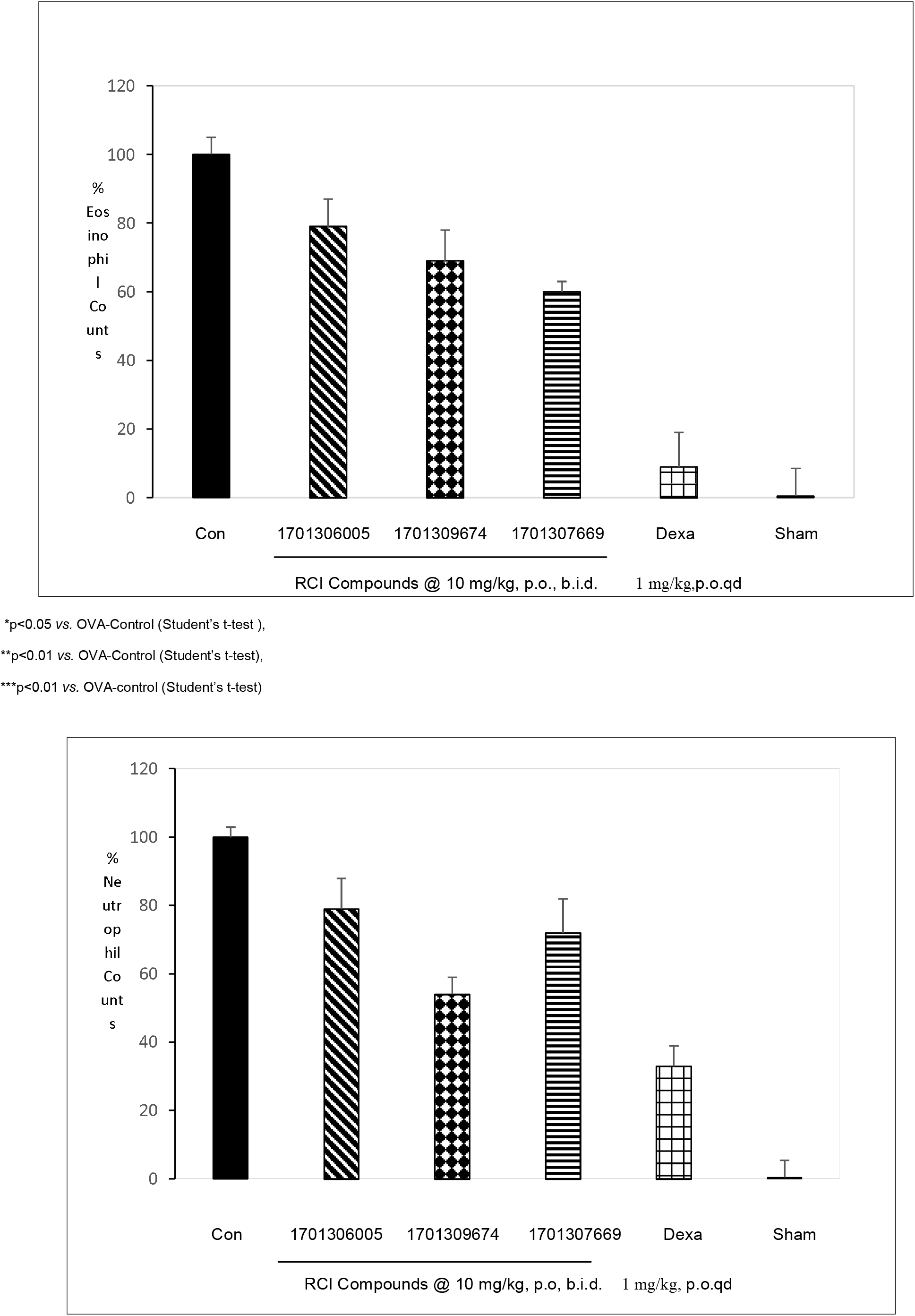
Evaluation of RCI compounds in HDM+CFA-induced airway inflammation model in BALB/c mice at 10 mg/kg, p.o. to monitor a) change in eosinophil and b) neutrophil counts.

**Figure-10.**
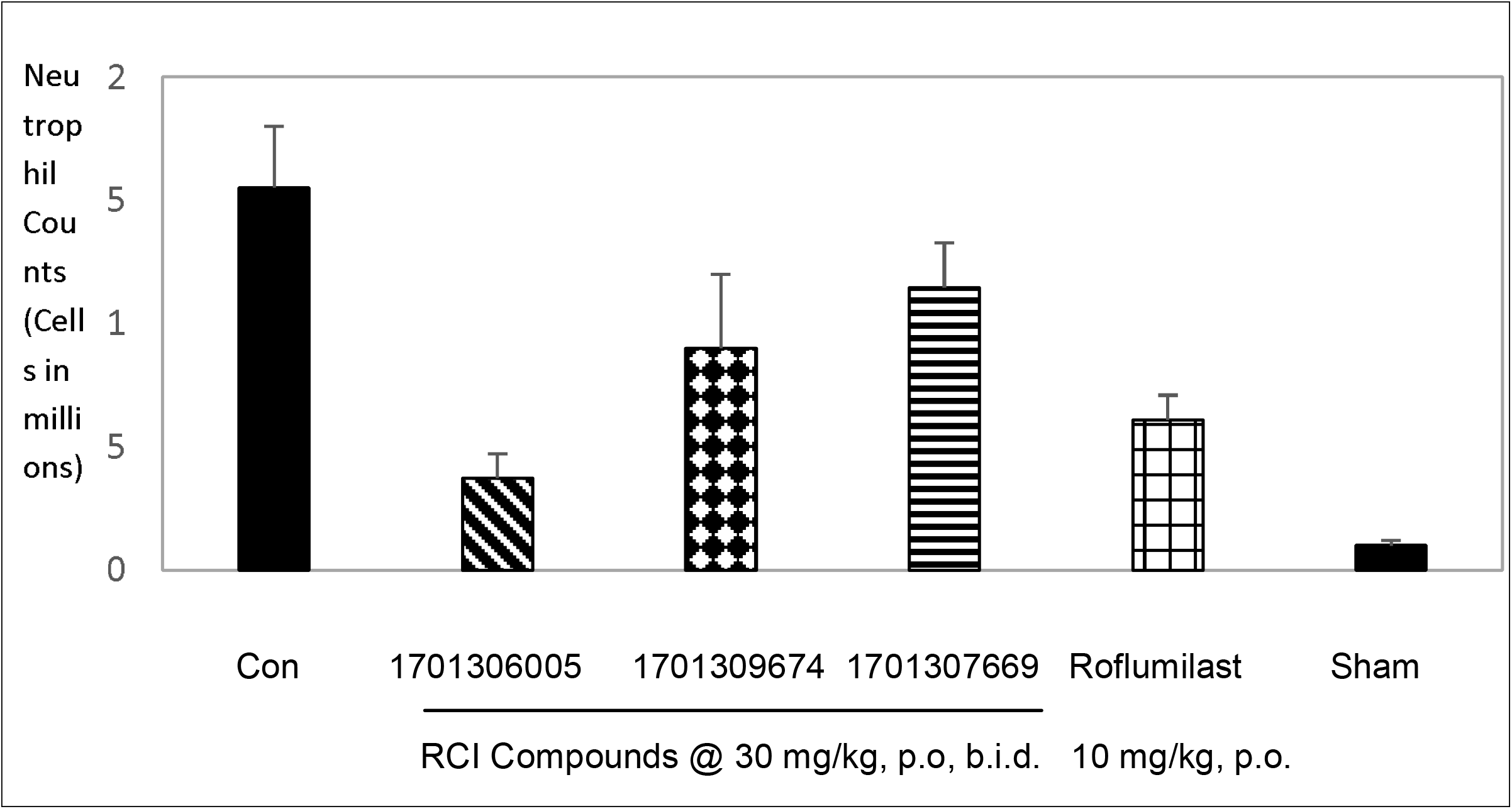
Evaluation of RCI compounds in LPS induced neutrophilia model in BALB/c mice at 30 mg/kg, p.o. to monitor the change in neutrophil counts as compared to vehicle control.

## 4. DISCUSSION

Asthma is a progressive disease and it has been reported that the disease severity is associated with an impairment in the biosynthesis of pro-resolving mediators in these patients. We hypothesized that the pro-resolving properties of FPR2 agonists, may suit well the chronic, non-resolving inflammatory conditions of the airways in asthma, and the development of selective FPR2 agonists seems to be an interesting approach to address the unmet medical need in patients with moderate to severe asthma.

We conducted structure-activity-relationship studies to identify potent FPR2 agonists which were selective for FPR1 and also possessed acceptable ADME and PK profile. Several molecules in the imidazole and benzimidazole series were identified as possessing potent FPR2 activity. These molecules were rank-ordered based on their in vitro profile and ADME properties (Table-1). RCI-1701306005, RCI-1701309674, and RCI-1701307669 were shortlisted further for detailed biological profiling. Shortlisted molecules showed low nM potencies for FPR2 agonism and >10,000 fold selectivity over FPR1 in the Ca^2+^ release assay and no activity in CHO-WT cells (Table-2). Selective FPR2 agonists in our study have demonstrated a time-dependent increase in the phosphorylation of extracellular signal-regulated kinase-1/2 (Erk1/2) in CHO cells overexpressing FPR2, as reported for FPR2 mediated response in some other studies also (Qin et al., 2017, Cattaneo et al., 2019; Smirnova et al., 2010). RCI compounds demonstrated a time-dependent increase in the phosphorylation of extracellular signal-regulated kinase-1/2 (Erk1/2) in CHO cells overexpressing FPR2 (Fig.3). PI3 kinase being an upstream member of the Erk pathway, we also tried to understand the involvement of PI3 kinase in FPR2 mediated functional response. Our data suggest that FPR2 agonists phosphorylate Erk1/2 in a PI3 kinase-independent manner. This further increases the scope of our study to identify the missing links of PI3 kinase-independent activation of Erk1/2 by FPR2 agonists. Since 15d-PGJ2 has been reported to be a mediator of resolution of inflammation by activating MEK/ERK pathway (Cash et al., 2014, Marcone et al., 2016; Oliva et al., 2003; Takeda et al., 2001), we wanted to know whether the selective FPR2 compounds in our study have any effect on 15d-PGJ2 release, which may be eventually leading to Erk1/2 phosphorylation directly or indirectly. FPR2 agonists have also been reported to inhibit COX2 in literature (Songjuan et al., 2019; Lan et al., 2021). Literature reports also suggest that COX2 inhibition leads to an increase in the levels of 15d-PGJ2 and mediate resolution (Hirotaka et al., 2002). We therefore explored the ability of FPR2 agonists for 15d-PGJ2 release to demonstrate the pro-resolution potential of FPR2 agonists in cell based assays. Our data suggest that RCI molecules increase the levels of 15-D PGJ2 in CHO-FPR2 cells (Fig.4). Since neutrophil migration is involved in the pathogenesis of bronchial asthma, we also tested these molecules to assess their effect on neutrophil migration. We used differentiated HL-60 cells, which have been previously (Millius et al., 2010) reported possessing neutrophil-like features. When RCI molecules were tested in the migration assay using HL-60 cells, they showed potent activity (Fig.5). FPR2 mRNA expression in HL-60 cells was confirmed by the real-time PCR experiment.

**Table-2.**

| Test Compound | Calcium release assay (EC<sub>50</sub> in nM) |  |  |  |
| --- | --- | --- | --- | --- |
|  | hFPR2-CHO | mFPR2-CHO | hFPR1-CHO | CHO WT |
| **RCI-1701306005** | 16.09±2.32 | 42.15±3.21 | >10000 | >10000 |
| **RCI-1701309674** | 3.48±0.41 | 3.34±0.72 | >10000 | >10000 |
| **RCI-1701307669** | 4.01±0.33 | 2.52±0.24 | >10000 | >10000 |
| **W peptide** | 0.56±0.22 | 0.015±0.01 | 1.27±0.23 | >10000 |

**2b-**Ca<sup>2+</sup> release assay in presence of FPR2 antagonist
| Compound | Calcium release assay in hFPR2-CHO cells (EC<sub>50</sub> in nM) |
| --- | --- |
| **W peptide** | 0.56±0.2 |
| **W peptide + WRW4 (10µM)** | >10000 |
| **RCI-1701307669** | 4.01±0.33 |
| **RCI-1701307669 + WRW4(10µM)** | >10000 |

**Table 3.** *In vivo* Pharmaco-kinetic data of RCI compounds (n=2)

| Parameters | C<sub>max</sub> |  | AUC |  | F% |
| --- | --- | --- | --- | --- | --- |
| Dose (mg/kg, p.o) | 100 | 10 | 100 | 10 | 10 |
| RCI-1701306005 | 7.9 | 1.7 | 27.3 | 1.6 | 53 |
| RCI-1701309674 | 7.8 | 1.7 | 82.5 | 2.8 | 57 |
| RCI-1701307669 | 9.4 | 1.1 | 122.1 | 3.7 | 78 |

RCI compounds with impressive *in vitro* biological profile and selectivity over FPR1 were evaluated further for their potential in mitigating the non-resolving inflammation in mouse models of moderate to severe asthma. Shortlisted; RCI-1701306005, RCI-1701309674, and RCI-1701307669 inhibited OVA-induced airway eosinophil influx in mouse, in a dose-dependent manner with ID50 values of 11.8, 10.7, and 14.7 mg/kg (bid, p.o.) respectively (Fig.7). Further to this, we evaluated RCI compounds in another two more challenging models of asthma using HDM and HDM+CFA. HDM is an environmental allergen that leads to chronic airway inflammation as well as airway hyper-responsiveness. It is therefore considered a risk factor for persistent asthma in human subjects. Interestingly, RCI compounds showed significant inhibition of eosinophil influx in the BALF of these mice at 10 and 30 mg/kg and moderate inhibition of airway hyper-responsiveness at 30 mg/kg in the mouse HDM model (Fig.8). Besides, these compounds showed inhibition of eosinophils and neutrophils counts in the bronchoalveolar lavage of mice in the HDM+CFA model of increased airway resistance (Fig.9), which is even a more challenging model. Potent *in vivo* efficacy demonstrated by RCI compounds in three distinct asthma models by us confirms the anti-asthmatic potential of FPR2 agonists. We, therefore, propose herewith that the development of selective FPR2 receptor agonists can be a potential approach for the management of moderate to severe asthma.

## ACKNOWLEDGEMENTS

We acknowledge Dr. Ashok Patra, Director, Imgenex India for discussions in the generation of FPR2 and FPR1 stable cell lines.

## CONFLICT OF INTEREST

All the authors declare that they have no conflicts of interest.

## Notes

### Competing Interest Statement

The authors have declared no competing interest.

